# Establishment and characterization of a cell line and patient-derived xenograft (PDX) from peritoneal metastasis of low-grade serous ovarian carcinoma

**DOI:** 10.1101/784959

**Authors:** Elien De Thaye, Koen Van de Vijver, Joni Van der Meulen, Joachim Taminau, Glenn Wagemans, Hannelore Denys, Jo Van Dorpe, Geert Berx, Wim Ceelen, Jan Van Bocxlaer, Olivier De Wever

**Author notes:** **Correspondence to:** Olivier De Wever, Laboratory of Experimental Cancer Research, Department of Radiation Oncology and Experimental Cancer Research, Faculty of Medicine and Health Sciences, Ghent University, Corneel Heymanslaan 10, B-9000 Ghent, Belgium.

## Abstract

Peritoneal spread indicates poor prognosis in patients with serous ovarian carcinoma (SOC) and is generally treated by surgical cytoreduction and chemotherapy. Novel treatment options are urgently needed to improve patient outcome. Clinically relevant cell lines and patient-derived xenograft (PDX) models are of critical importance to therapeutic regimen evaluation. Here, a PDX model was established by orthotopic engraftment, subperitoneal tumor slurry injection, of low-grade SOC resulting in an early-stage transplantable peritoneal metastasis (PM)-PDX model. Histology confirmed the micropapillary and cribriform growth pattern with intraluminal tumor budding and positivity for PAX8 and WT1. PM-PDX dissociated cells show an epithelial morphotype with a 42h doubling time and 40% colony forming efficiency, they are insensitive to estrogen signaling, low sensitive to platinum derivatives and highly sensitive to paclitaxel (IC50: 6.3 ± 2.2 nM, mean ± SE). The patient primary tumor, PM, PM-PDX and derived cell line all show a *KRAS* c.35G>T (p.(Gly12Val)) mutation and show sensitivity to the MEK inhibitor trametinib in vitro (IC50: 7.2 ± 0.5 nM, mean ± SE) and in the PM mouse model. These preclinical models closely reflecting patient tumors are useful to further elucidate LGSOC disease progression, therapy response and resistance mechanisms.

## Background

Ovarian cancer, the deadliest gynecological cancer, is the eight most frequently diagnosed cancer and ranks as the eight leading cause of cancer death in women, with an estimated 300 000 new cases and 185 000 deaths in 2018 worldwide (1). Ovarian cancer is a very heterogeneous disease. The most common type is high-grade serous ovarian carcinoma (HGSOC) which account for 70-75% of all ovarian malignancies (2). The vast majority are characterized by *TP53* mutations and lack mutations of *KRAS, BRAF* or *ERBB2*. Low-grade serous ovarian carcinoma (LGSOC) accounts for less then 5% of all ovarian serous carcinomas, other epithelial ovarian cancer types are endometrioid (8-10%), clear cell (8%), seromucinous (3%), mucinous (3%) and Brenner (1%) tumors (3). LGSOC is characterized by mutations of the *KRAS, BRAF* or *ERBB2* genes, in which approximately two thirds of tumors have a mutually exclusive mutation in one of these genes (4). *KRAS, BRAF* and *ERBB2* are upstream activators of the mitogen-activated protein kinase (MAPK) pathway, leading to cellular proliferation. As both types of cancer are associated with vague symptoms in early stages, the majority of patients present with advanced-stage disease (5). The presence of peritoneal carcinomatosis, which results from intra-abdominal metastases, is associated with the late presentation of the disease. Treatment difficulties of peritoneal metastases and the possible recurrences do both contribute to a poor prognosis of this cancer (6). Given the high relapse rate and poor prognosis of this disease, interest increases in the development of new treatment approaches (7). Therapeutic management of ovarian cancer has traditionally been based on a combination of surgery and platinum-/taxane-based chemotherapy (6). However, LGSOC is not as responsive to platinum-/taxane-based chemotherapy as HGSOC. Although a clear involvement of the MAPK pathway in the disease is demonstrated, a phase 3 study using the MEK inhibitor binimetinib showed mid-term discontinuation, most probably due to escape mechanisms leading to lack of treatment efficacy (8).

In every aspect of translational cancer research, from the biological aspects of the disease to the development of new treatments, the use of preclinical models is a key component. In recent years, there has been an increasing interest in the application of organoids and patient-derived xenografts (PDXs) because of their high potential as an essential tool for personalized medicine (9–11). The process of generating PDXs (also known as tumorgraft models) is based on the transfer of fresh tumor tissue (primary or metastatic) from the patient directly to an immunocompromised mouse (12).

Depending on the cancer type, pretreatment, amount of tissue available, molecular properties etc., the success rate of the PDX will vary (13). The organ environment can affect tumor engraftment, highlighting the role of the site of implantation. Traditionally the tumor fragment is implanted into an area unrelated to the original tumor site, which is considered a heterotopic implantation (generally subcutaneous). On the other hand, tumor xenografts can also grow orthotopically into the corresponding anatomic region but their use is often hindered by a need for a high level of technical skills, time and cost (14). For some cancers, such as colorectal, breast, lung, pancreatic, head and neck, melanoma, gastric, ovarian, prostate and renal cancer, methodologies for PDX establishment and characterization are already described in literature with engraftment rates ranging from 9 to 90% of success (13, 15).

In this work, for the first time, an orthotopic PDX model, based on a subperitoneal tumor slurry injection, and cancer cell line from a peritoneal metastasis of LGSOC were established. This model showed a *KRAS* mutation and sensitivity to the MEK inhibitor trametinib demonstrating its clinical relevance to study treatment responsiveness and resistance mechanisms.

## Methods

### Establishment of peritoneal metastasis (PM)-PDX models

Fresh peritoneal tissue specimens from 10 consenting patients with metastatic serous epithelial ovarian cancer (FIGO stage III or IV) were collected at the time of debulking surgery at Ghent University Hospital, Belgium. Nine patients were diagnosed with HGSOC and one with LGSOC. The study protocol was approved by the institutional review board of the Ghent University Hospital and the trial is registered as ClinicalTrial.gov NCT02567253 with EudraCT number 2015-000418-23. Samples were processed to a tumor slurry and injected in SCID/Beige mice within 75 minutes after removal from the patient. Tumors were minced in limiting volumes of RPMI 1640 media (Life Technologies, Ghent, Belgium), supplemented with 100 U/ml penicillin and 100 μg/ml streptomycin (Life Technologies, Ghent, Belgium). After a centrifugation step at 1500×g for 3 minutes, the upper culture medium was removed and tumor tissue was suspended in 1:1 Matrigel (Corning, The Netherlands). Further, a laparotomy was performed and 50 μL of the tumor suspension using a 19G needle was injected bilateral subperitoneally in three female 4 to 5 week old SCID/Beige (C.B-17/IcrHsd-*Prkdc^scid^Lyst^bg-J^*) mice (Envigo, The Netherlands). Animal studies were conducted in accordance with the local committee on the Ethics of Animal Experiments (Ghent University Hospital, Ghent, Belgium [ECD 15/28]). Cryopreserved tumors were minced and stored 1:1 in freezing media (90% FBS/10% DMSO) at −80°C and then in liquid nitrogen indefinitely.

### Tissue processing and immunohistochemistry

Tissues collected from mice or patients were fixed overnight in neutral buffered 10% formalin solution (Sigma-Aldrich, Belgium) and processed in the lab (H&E staining) or in the tissue core facility at Ghent University Hospital (immunohistochemistry).

### *In vivo* imaging

Transparent ultrasound transmission Polaris II gel (Ondes & Rayons Medical, France) was applied to bare skin and a MicroScan™ MS550D (22–55 MHz, VisualSonics Inc., Canada) transducer with the Vevo^®^ 2100 imaging system (VisualSonics Inc., Canada) was used to analyse the tumor cross-sectional area in Vevo LAB 1.7.1 (VisualSonics Inc., Canada).

### Establishment of tumor-derived cell lines

To establish cell lines derived from the peritoneal metastasis and a PM-PDX-model, tumor samples were cut into pieces of 2-4 mm and suspended using the tumor dissociation protocol with the GentleMacs^®^ dissociator (Miltenyi Biotec GmbH, Germany). The cell suspension was applied to a cell strainer (70 μm, Corning, The Netherlands), centrifuged at 300×g for 7 minutes and after aspiration of the supernatant resuspended in complete EMEM supplemented with 10% fetal bovine serum, 100 U/ml penicillin and 100 μg/ml streptomycin (Life Technologies, Ghent, Belgium). The first weeks, cells were maintained in a 6-well plate (Novolab, Belgium) before culturing in a T25 falcon at 37°C and 5% CO_2_ in air. The cell culture was monthly tested for Mycoplasma by using MycoAlert Plus Kit (Lonza, Basel, Switzerland).

### *KRAS* mutation analysis

Exons 2, 3 and 4 of the *KRAS, NRAS* and *HRAS* genes and exon 15 of the *BRAF* gene were analysed using a PCR-based enrichment strategy followed by library preparation and MiSeq sequencing. In brief, DNA was extracted using the QIAamp DNA Blood mini kit (Qiagen) for cell culture samples or using the QIAamp DNA FFPE Tissue kit and deparaffinisation solution (Qiagen) for formalin-fixed paraffin-embedded (FFPE) slices. The DNA concentration was measured by use of the Trinean Dropsense96 UV/VIS droplet reader (Trinean) or with Qubit (Thermofisher). For the PCR, the KAPA2G Robust mastermix was used together with 0.5 μM primers and 10 ng of DNA template in a 30 μl reaction volume. The PCR protocol consists of 5 min at 95°C, 50 cycles (30 sec at 95°C, 45 sec at 60°C and 45 sec at 72°C) and 1 min at 72°C. Library preparation made use of the Nextera XT kit (Illumina) and massive parallel sequencing was performed on MiSeq (Illumina) (16). All PCR and massive parallel sequencing reactions were performed in duplicate. Data-analysis was performed by use of the commercial software package CLC bio Genomics Workbench v9 (Qiagen).

### Luciferase-EGFP transduction

293T cells were cultured in DMEM (41965039, ThermoFisher) with 10% Fetal Calf Serum (FCS) (Sigma-Aldrich, St Louis, MO, USA) and 2 mM L-Glutamine (BE17-605F, Lonza) and transfected with lentiviral envelope plasmid pMD2.G, packaging plasmid psPAX2 and lentiviral expression plasmid pLenti6-LUC2CP-EGFP-Blast. The medium was removed and replaced with fresh medium 8 hours post transfection. The virus was harvested 48 hours post transfection and filtered through a 0.45 μm PES filter (Merck-Millipore, Burlington, Massachusetts, USA). PM-PDX derived cells were cultured in complete EMEM until a density of approximately 60% was reached. The medium was removed and replaced by pLenti6-LUC2CP-EGFP-Blast virus containing medium for 24 hours. Cells expressing the construct were selected after addition of 2.5 μg/ml Blasticidin S (R21001, ThermoFisher) to the medium. After 10 days the cells expressing the LUC2CP-EGFP fusion protein were sorted with the BD FACSAria III cell sorter.

### Clonogenic assay

500 PM-PDX-derived cells were seeded in different T25 cell culture flasks and immediately treated with 15, 150 or 1500 pg/ml estrogen or 1, 10 or 100 nM trametinib, selumetinib or fulvestrant. Control conditions were 0.1% DMSO or stripped medium for the estrogen experiment. Cells were incubated during 8 days in the presence of the drug (3 T25 flasks/condition) and effectiveness of all agents was determined by staining the colonies using crystal violet as an endpoint measurement.

### IncuCyte ZOOM monitored studies

Real-time monitoring of cell confluency was performed using the IncuCyte ZOOM System (Essen Bioscience, Hertfordshire, UK) according to the manufacturer’s guidelines. For cell confluency monitoring, cells were seeded in 96-well clear-bottom Corning^®^ Costar^®^ cell culture plates at 2 000 cells per well (100 μl/well) and allowed to adhere 24 hours at 37°C and 5% CO_2_ in air. Subsequently, cells were exposed to the drugs in complete EMEM medium and microscopic images (4 images/well) were taken every two hours for the duration of the experiment. All images were analysed and cell confluency was deduced using IncuCyte software. Each condition was performed in, at least, four fold. Chemotaxis cell migration was studied for SK-OV-3 luc IP1 and PM-LGSOC-01 cells using the IncuCyte™ ClearView 96-Well Cell Migration plate coated with 1% Matrigel (Corning, The Netherlands) in 0% FBS EMEM medium. 3 000 cells/well were seeded (60 μL volume) with 0.1% FBS to the top and 200 μL 10% FBS to the bottom. Cell migration was followed using the phase contrast cell confluency monitoring.

### Cell lysates and western blotting

Proteins were extracted from the cells using the Laemmli lysis buffer (0.125 M Tris-HCl, 10% glycerol, 2.3% sodium dodecyl sulfate (SDS), pH 6.8). After an ultrasonication step, cell lysates were suspended in reducing sample buffer (1 M Tris-HCl, 30% glycerol, 6% SDS, 3% β-mercaptoethanol, 0.005% bromophenol blue, pH 6.8) and boiled for 5 minutes at 95°C. 20 μg proteins of the cell line were exposed to a 10% SDS-PAGE gel and transferred to nitrocellulose membranes (Bio-Rad, Hercules, CA, USA). After blocking the membranes using 5% non-fat milk or bovine serum albumin (BSA) in phosphate-buffered saline (PBS) with 0.5% Tween 20 (Sigma-Aldrich, Belgium), the membranes were incubated overnight at 4°C with the primary antibodies (Table 1). After washing the membrane, incubation with HRP-conjugated secondary antibody was performed at room temperature for 1 hour. WesternBright Quantum HRP substrate (Advansta, Menlo Park, CA, USA) was added to the membranes to capture the luminescent signal using the Proxima 2850 Imager (IsoGen Life Sciences, De Meern, The Netherlands). Equal loading of samples was verified by primary monoclonal mouse anti-GAPDH antibodies (clone GAPDH-71.1, Sigma-Aldrich, Belgium).

**Table 1.** Primary antibodies used in western blot (WB)

| Primary antibodies | Source | WB |
| --- | --- | --- |
| Rabbit anti-α-catenin | Sigma Aldrich | 1:3000 |
| Mouse anti-α-smooth muscle actin (SMA) | Sigma | 1:1000 |
| Rabbit anti-β-catenin | Sigma Aldrich | 1:3000 |
| Mouse anti-E-cadherin | Invitrogen | 1:1000 |
| Mouse anti-estrogen receptor (ER)-α | Abcam | 1:1000 |
| Mouse anti-pan-cadherin | Sigma Aldrich | 1:1000 |
| Mouse anti-pan-cytokeratin | Sigma Aldrich | 1:1000 |
| Mouse anti-p53 | Sigma Aldrich | 1:1000 |
| Mouse anti-vimentin | Sigma Aldrich | 1:1000 |
| Mouse anti-GAPDH | Sigma | 1:1000 |
| Rabbit anti-phospho-p44/42 MAPK (ERK1/2) | Cell signalling Technology | 1:3000 |
| Rabbit anti-p44/42 MAPK (ERK1/2) | Cell signalling Technology | 1:3000 |

### *In vivo* PM-PDX-derived cell line model and animal study

Female 4-week-old SCID/Beige (C.B-17/IcrHsd-*Prkdc^scid^Lyst^bg-J^*) mice (Envigo, The Netherlands) were treated daily via oral gavage with vehicle (0.5% methylcellulose and 0.2% Tween 80 in water, n = 6) or trametinib (0.3 mg/kg/day, n = 6). Mice were treated starting 1 week after intraperitoneal injection of 1×10^6 Luciferase-EGFP expressing PM-PDX-derived cells (1:1 serum free EMEM medium:Matrigel (Corning, The Netherlands)). After 5 weeks of oral treatment, mice were sacrificed. Tumour development was assessed by weekly bioluminescence imaging until six weeks after cell injection. In order to measure bioluminescent signals, mice were given an intraperitoneal injection of 100 μL Xenolight D-luciferin (K+ Salt, Perkin Elmer, Belgium) in DPBS (without Ca^2+^ and Mg^2+^, 150 mg/kg mouse) and were anaesthetized with isoflurane (5% in oxygen for induction and 1.5% in oxygen for maintenance, IsoFlo, Abbott, Belgium). Imaging was initiated 15 minutes after injection using the IVIS Lumina II (Caliper Life Sciences). Exposure times were set automatically.

### Statistical analysis

Results obtained with the colony formation assay were analysed using one-way ANOVA with Tukey’s multiple comparisons test using Graphpad Prism 7 (GraphPad Software, USA). Using R Studio (17), Mann-Whitney U test was used to compare differences in relative total flux between groups in the *in vivo* experiment. Statistical tests were two-sided and p-value below 0.05 was considered statistically relevant. In figures, * represents p-value ≤ 0.05, ** p-value ≤ 0.01 and *** p-value ≤ 0.001.

## Results

### A low grade serous ovarian carcinoma (LGSOC) peritoneal metastasis (PM)-PDX model

Figure 1A illustrates the establishment of the LGSOC PM-PDX model. Based on the observed increase in high-density signal from the ultrasound imaging (Figure 1B), it was decided to passage the tumor tissue to a new group of acceptor mice 46 days after injection. This second passage was monitored over 3 months but no changes in high-density ultrasound signal was demonstrated. At day 146 post implantation, a macroscopically blister-like appearance of the tumor area was observed which was formalin fixed and processed for (immuno)histology. H&E revealed the micropapillary and cribriform growth pattern typical for LGSOC surrounded by a large mass of stroma in the first PDX passage. In the second PDX passage this micropapillary pattern dominated the tumor area showing a single layer of epithelium forming a large lumen (Figure 1C). This micropapillary pattern was further characterized by intraluminal tumor budding. Immunohistochemical stainings for paired box gene 8 (PAX8), WT1, tumor suppressor protein p53, estrogen receptor (ER) and progesterone receptor (PR) further confirmed the typical characteristics of LGSOC (Figure 1D).

**Figure 1.**
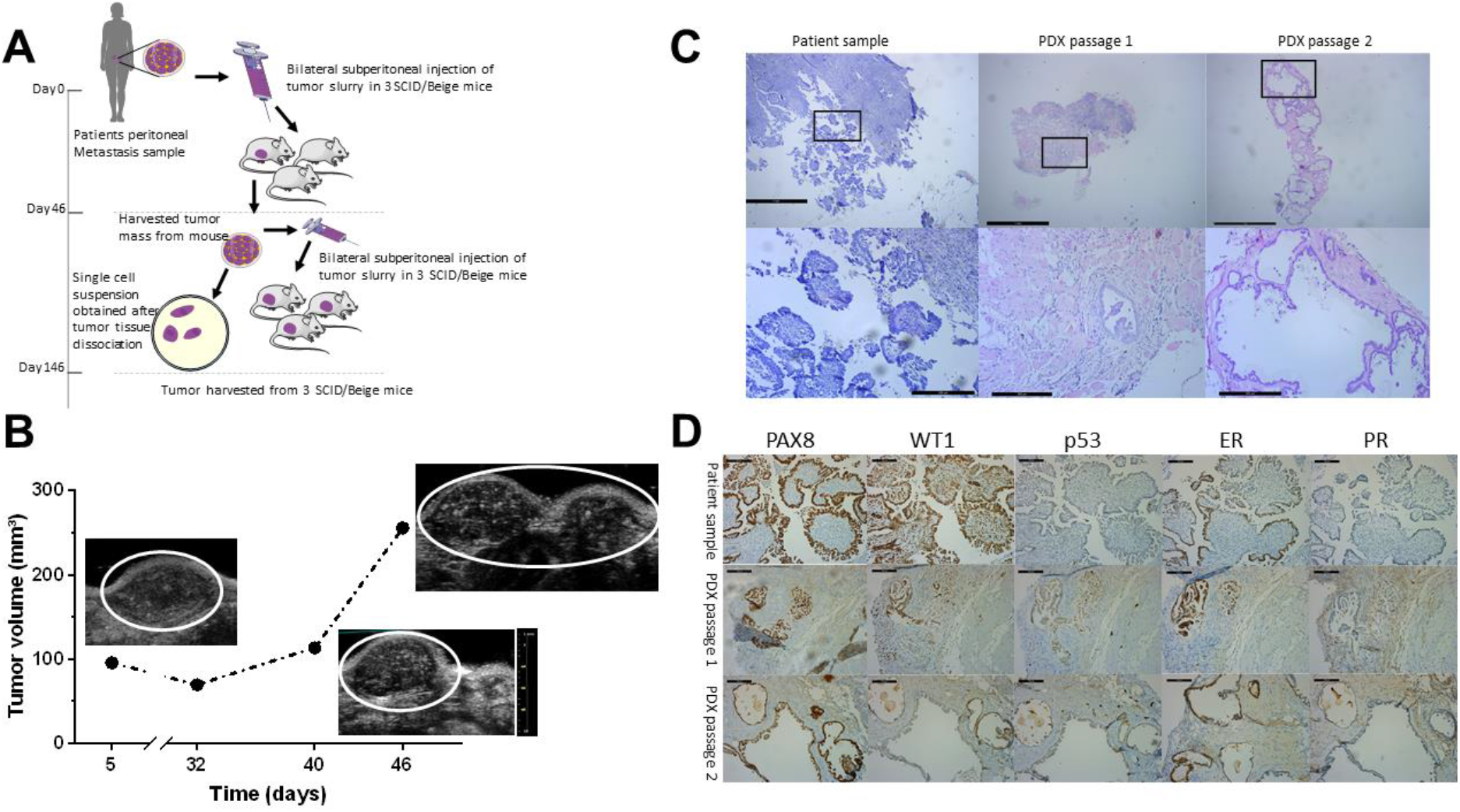
Establishment of the PM-PDX model. (A) Schematic representation of the protocol for PM-PDX model establishment. Freshly human peritoneal metastasis samples, originating from serous ovarian cancer, were collected and subperitoneally injected as a tumor slurry in SCID/Beige mice. The tumor is harvested once it is ready for passaging, tumor tissue is collected and prepared for subperitoneal injection in a new group of mice or processed into a single cell suspension. (B) Assessment of tumor volume over time using ultrasound imaging. (C) Tumor section slides were stained for H&E to compare histology of the PDX tumors with the corresponding patient metastasis. The lower row shows a close-up of the area within the black rectangle. Scale bars represent 1 mm for the upper row and 200 μm for the lower row. (D) Comparative study of tumor sections stained for PAX8, WT1, p53, ER and PR, as indicated. Scale bars represent 100 μm.

### Characterization of tumor-derived cell lines

Primary culture from single cell suspension of patient-derived peritoneal metastasis resulted in spread-polarized cells that typically showed signs of senescence characterized by a larger surface area and stress fibers (Figure 2A). These cultures showed a mixed expression of cytoskeletal proteins alfa-smooth muscle actin and cytokeratin and cell-cell adhesion molecules epithelial (E-) and neural (N-) cadherin, and most likely can be considered as mixed mesothelial-fibroblast cultures. In contrast, primary culture starting from tissue of the first passage PM-PDX model resulted in typical epithelial cells with cobblestone organization with strong cell-cell adhesion that showed colony growth. The first 5 to 8 initial subcultures showed no constant timing (among 2 to 3 weeks), the period in which cell proliferation was slow and unable to cover the entire culture flask surface. After this period, cell proliferation became quicker and in vitro passages for the maintenance of cell culture became regular (every week). The cell culture, named as PM-LGSOC-01, has been in continuous culture for >30 months and >100 in vitro passages (Figure 2A). PM-LGSOC-01 cells had a doubling time of 42 hours at passage 5 that was reduced to 23 hours at passage 22 and later passages. Table 2 summarizes the main findings regarding STR analysis. Comparison of STR profiles between PM-LGSOC-01 and other human cell lines did not match evaluation values greater than 0.82, confirming the uniqueness of PM-LGSOC-01 cell line. Results of western blotting (Figure 2B) illustrate a stable expression of cytoskeletal and cell-cell adhesion proteins over a wide range of passage numbers. Despite the presence of E-cadherin and its associated cytoplasmic catenins the PM-LGSOC-01 cells form aggregates but do not show compact spheres within 48h in contrast to positive controls used for compact sphere formation (Figure 2C). Chemotactic migration of PM-LGSOC-01 to a 10% FBS gradient was limited in contrast to SK-OV-3 luc IP1 cells characteristically used as a migratory ovarian cancer cell line (Figure 2D).

**Figure 2.**
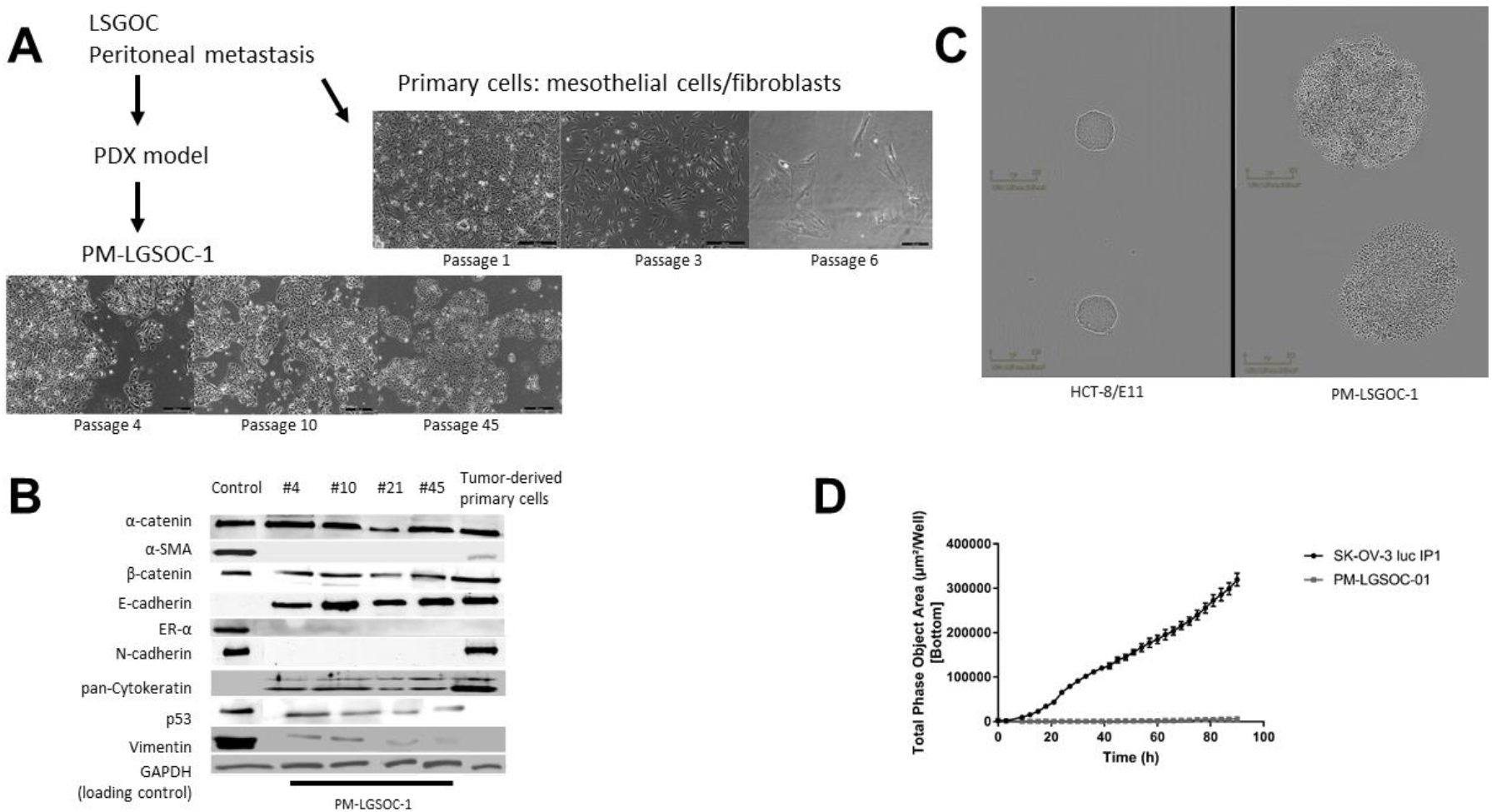
Characterization of tumor-derived cell lines. (A) Morphology of tumor-derived primary cells, directly derived from patient material or after one passage in mice. Scale bars represent 500 μm for the tumor-derived primary cells and 200 μm for the PM-LGSOC-01 cells. (B) Immunoblotting results for different in vitro passages of the PM-LGSOC-01 cell line and the tumor-derived primary cells. CT5.3hTERT cells were used as reference and MCF-7/AZ cells were used as a reference for ER-α expression levels. GAPDH was used as the loading control. (C) Evaluation of the aggregation activity of the PM-LGSOC-01 cells using IncuCyte technology. HCT-8/E11 cells were included as positive controls for compact sphere formation. Upper and lower panel indicate two separate experiments. Scale bars represent 300 μm. (D) Real-time monitoring of migration activity of SK-OV-3 luc IP1 cells and the PM-LGSOC-01 cells using the IncuCyte technology. The evaluation was performed using 0.1% FBS in culture medium on top and 10% FBS in culture medium at the bottom. Mean ± SE of six technical replicates is shown.

**Table 2.**
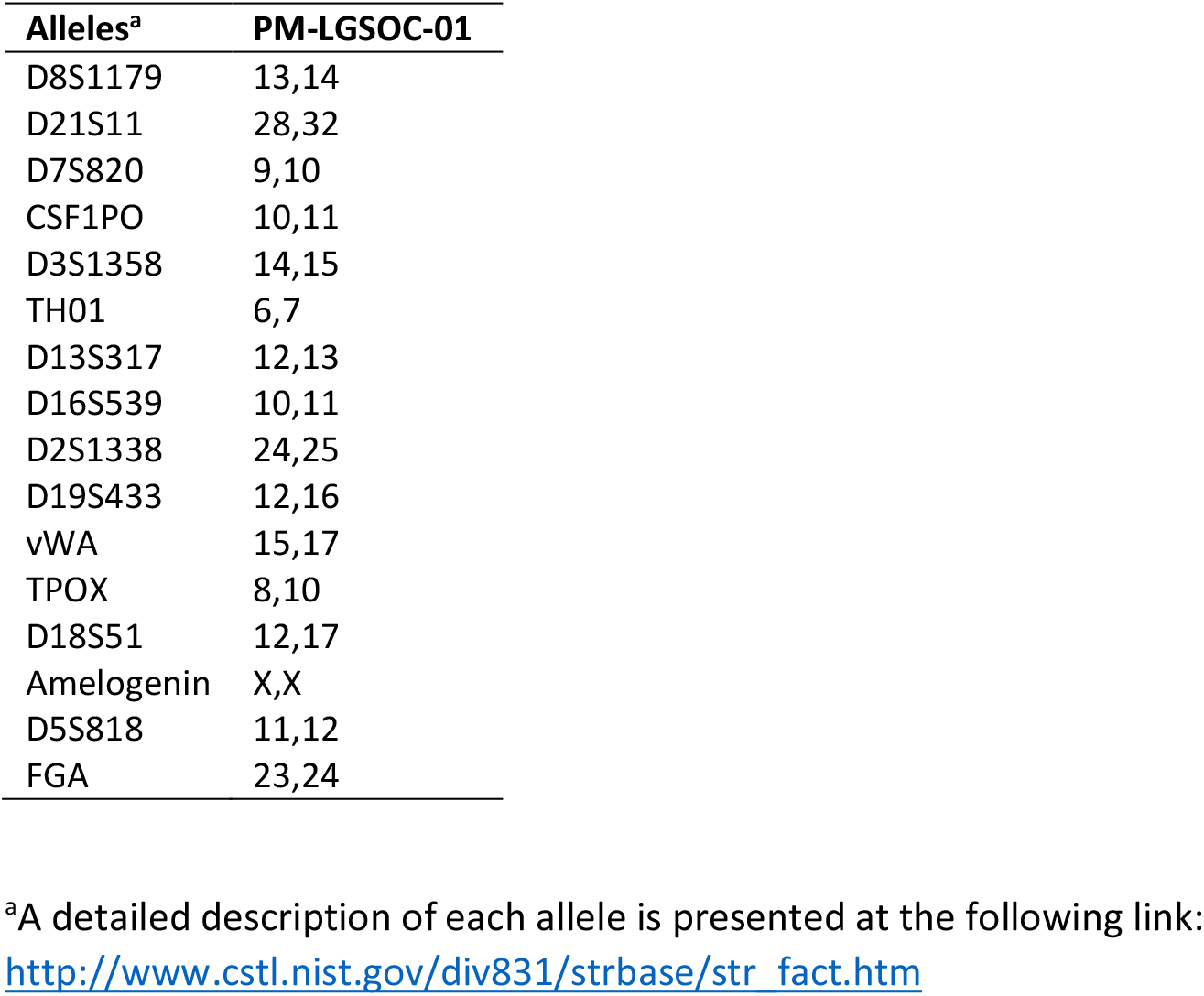
STR profile for the PM-LGSOC-01 cell line

| Alleles<sup>a</sup> | PM-LGSOC-01 |
| --- | --- |
| D8S1179 | 13,14 |
| D21S11 | 28,32 |
| D7S820 | 9,10 |
| CSF1PO | 10,11 |
| D3S1358 | 14,15 |
| TH01 | 6,7 |
| D13S317 | 12,13 |
| D16S539 | 10,11 |
| D2S1338 | 24,25 |
| D19S433 | 12,16 |
| vWA | 15,17 |
| TPOX | 8,10 |
| D18S51 | 12,17 |
| Amelogenin | X,X |
| D5S818 | 11,12 |
| FGA | 23,24 |

<sup>a</sup>A detailed description of each allele is presented at the following link:
http://www.cstl.nist.gov/div831/strbase/str_fact.htm

### *In vitro* effect of trametinib on *KRAS* mutated PM-LGSOC-01 cells

Evaluation for typical mutations of LGSOC found that the patient’s primary tumor and peritoneal metastasis, PDX passage 1 and the PM-LGSOC-01 cell line early and late passages (3, 32 and 72) and its luc-EGFP transduced PM-LGSOC variant all carried the *KRAS* c.35G>T (p.(Gly12Val)) mutation, as illustrated in Figure 3A. Due to the presence of this mutation, the efficacy of the MEK inhibitors trametinib and selumetinib was further investigated. Indeed, trametinib dose-dependently inhibits ERK phosphorylation and cell confluency with an IC50 of 7.2 ± 0.5 nM (mean ± SE) (Figure 3B). Selumetinib also affected cell confluency but only in higher molar concentrations. In agreement with the poor chemosensitivity of LGSOC only paclitaxel shows a sensitivity in the low nM range (IC50 of 6.3 ± 2.2 nM (mean ± SE)) in contrast to platinum based compounds with IC50 > 2 μM (Figure 3C). In agreement, the clonogenic assay confirmed the effect of trametinib and selumetinib on clone numbers (Figure 3C). Cell cycle analysis confirmed the impact of trametinib on cellular growth by stimulating a cell population into an increased G1 phase and decreased S and G2/M phase (Figure 3D).

**Figure 3.**
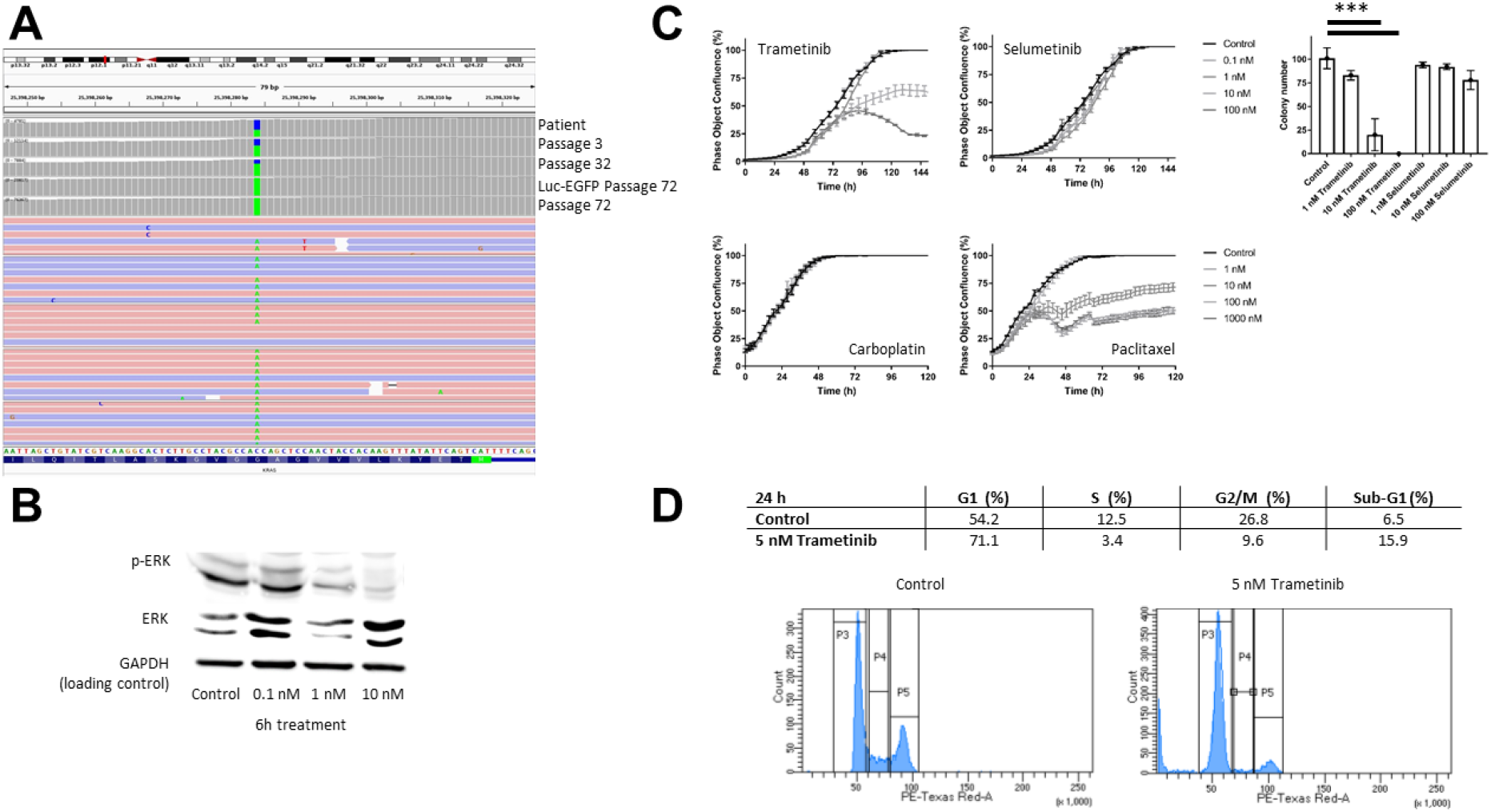
*In vitro* effect of trametinib on *KRAS* mutated PM-PDX derived cells. (A) *KRAS* c.35G>T (p.(Gly12Val)) mutation analysis at patient material, different in vitro passages of the PM-LGSOC-01 cell line (3, 32 and 72) and for the Luc-EGFP positive PM-LGSOC-01 cells. The colors blue and green indicate the fraction wildtype versus mutant, respectively. (B) Immunoblotting results for p-ERK and ERK of the PM-LGSOC-01 cell line treated with 0.1% DMSO (control) and trametinib at a concentration of 0.1, 1 and 10 nM for 6 hours. GAPDH was used as the loading control. (C) On the left, real-time analysis of PM-PDX derived cell confluency using IncuCyte technology. PM-LGSOC-01 cells were treated with 0.1% DMSO (control), trametinib, selumetinib, carboplatin and paclitaxel at concentrations of 0.1, 1, 10, 100 and 1000 nM. Mean ± SE of at least four technical replicates is shown. On the right, results on the clonogenicity assay. PM-LGSOC-01 cells were treated for 1 week with trametinib or selumetinib at a concentration of 1, 10 and 100 nM. Mean + SE of three technical replicates is shown. Statistical analysis was performed using one-way ANOVA at the α = 0.05 significance level. (D) Results of the cell cycle distribution analysis by flow cytometry. Quantitation of the sub-population fractions of the histograms. PM-LGSOC-01 cells were treated for 24 hours with 0.1% DMSO (control) or 5 nM trametinib. Many cells were blocked in the G0/G1 phase and a reduction in the S and G2/M phase was observed with increasing concentration of trametinib.

### Impact of trametinib in an *in vivo* peritoneal metastasis model of LGSOC

PM-LGSOC-01 cells were lentiviral transduced to obtain constitutive GFP- and Luciferase expression. These reporter cells were further used to create a peritoneal metastasis model from LGSOC in order to evaluate the effect of trametinib *in vivo*. Figure 4A illustrates the imaging data at different time points before and during the treatment period. In both groups, during the time course of the experiment no mice developed ascites. Animals received daily oral gavage based on vehicle or 0.3 mg/kg trametinib in a volume of 100 μL. Over time a clear increase in bioluminescence activity can be observed for the control group whereas a decrease in signal is observed in the trametinib treatment group. After 5 weeks of treatment animals were euthanised and relative total flux was significantly higher in the control group compared to the trametinib group (Figure 4B). On average a 4-fold increase in bioluminescent increase from the start of the experiments was observed for the control group whereas on average the bioluminescent signal decreased with about 30% in the trametinib group, relative to starting conditions. Figure 4C illustrates the histopathological (H&E) and immunohistochemical stainings (Ki67 and PAX8) representative for both the control and trametinib group. H&E shows nests of cells that organize into papillae surrounded by stroma characteristic of LGSOC. Ki67 labeling index was twice as high in the control group (30%) compared to the trametinib group (15%).

**Figure 4.**
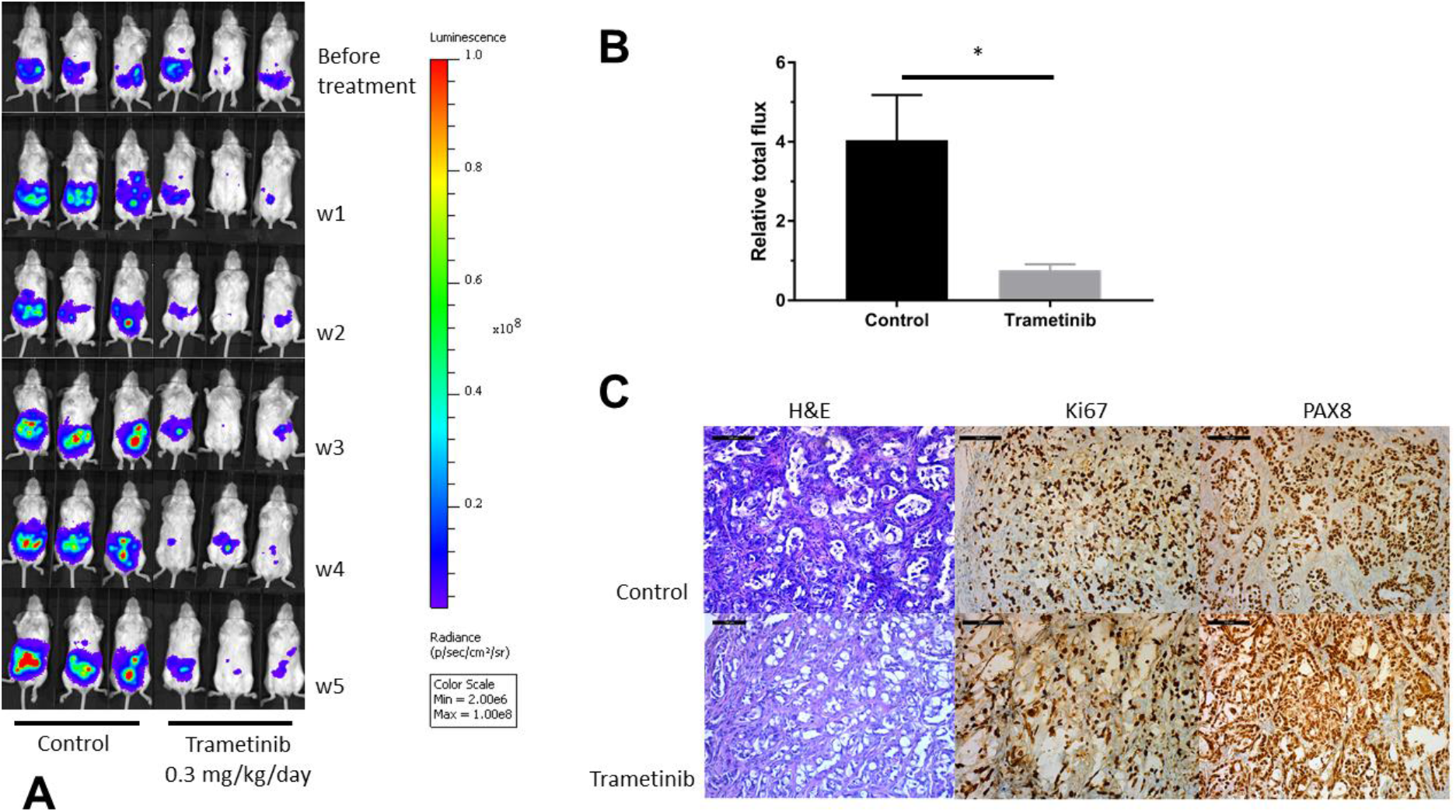
Impact of trametinib in an *in vivo* peritoneal metastasis model. (A) Monitoring of in vivo bioluminescence in SCID/Beige mice after intraperitoneal inoculation of Luciferase-EGFP positive PM-LGSOC-01 cells and treated daily with vehicle or trametinib (0.3 mg/kg/day) via oral gavage. (B) Bar plot indicating the increase in bioluminescent signal, detected after 5 weeks of daily treatment, corrected for the observed signal before therapy per individual mouse (relative total flux). Data represent mean + SE of five animals/group. (C) Histopathological (H&E) and immunohistochemical (Ki67 and PAX8) analysis of tumor sections representative for the control and treatment group. Scale bars represent 100 μm.

## Discussion

The heterogeneous nature of ovarian cancer makes it challenging to predict therapeutic responses in patients (18, 19). In this regard, preclinical models accurately mimicking biological properties of in vivo human tumors are of great value for efficient drug discovery (20). To date, preclinical research in LGSOC has been limited. The low frequency and slow growth rate of these tumors have challenged the development of cell lines and animal xenograft models. LGSOC cell lines are not available at the American Type Culture Collection (ATCC) and are only available at some research groups worldwide (21, 22). Kopper et al. (2019) established organoid lines in basement membrane extracts representing both LGSOC and HGSOC from primary tumor, ascites and peritoneal metastasis (11). The organoid lines allow subcutaneous transplantation and can be used in drug screening assays. Our approach was slightly different. A tumor slurry from peritoneal metastatic tissue of LGSOC was subperitoneally injected into an immunodeficient SCID/Beige mouse leading to tumor growth. From this early-stage PDX a tissue-culture substrate adherent cell line was established that showed long term in vitro expansion and enabled manipulation and functional analysis. We also confirmed the histological features of the early-stage PDX such as micropapillae surrounded by stroma in the first passage and marked architectural complexity in the second passage most probably due to anastomosis of micropapillae forming the elongated and branching structures. The genomic aberration characterized by *KRAS* mutation is consistent in the PM-PDX and PM-LGSOC-01 cell line. Biomarker expression, such as positive PAX8 and WT1 combined with a wildtype p53 is consistent in the primary tissue versus the PM-PDX and PM-LGSOC-01, even after extended passage. Ovarian PDXs are predominantly originating from HGSOC as a low take rate and long latency is often associated with other histological subtypes. However, in our case HGSOC patients were strongly pretreated by chemotherapy and characterized by necrotic areas and areas containing cancer cells with low mitotic activity making it less likely to establish a PDX model from PM of HGSOC patients. In contrast, the LGSOC patient did not receive neoadjuvant chemotherapy before surgery leading to more viable tumor tissue, easily forming an early-stage transplantable PDX and generated tissue-culture adherent PM-LGSOC-01 cell line, low sensitive to platinum derivatives. Other characteristics are clonogenicity and tumorigenicity, lack of serum-induced chemotactic migration and absence of compact sphere forming activity despite the presence of cell-cell adhesion molecule E-cadherin and its downstream catenins. PM-LGSOC-01 cell line allows genetic manipulation and easy in vivo monitoring of its luc-EGFP variant using bioluminescence imaging. The mouse passaging of PM tumor tissue was necessary to obtain a tissue-culture adherent cell line since cells cultured directly from patient PM tumor tissue ended into dominant growth of stromal cells such as fibroblasts and mesothelial cells that become senescent after further passaging.

Prior studies have reported that LGSOC tumors have a unique clinical, pathological and molecular profile compared to other ovarian cancers. LGSOC harbours *KRAS* mutations in 19 to 54.5% of the cases and lacks *TP53* mutations (23–28). With the focus on inhibiting KRAS signalling via downstream effector MEK, both allosterically active compounds trametinib and selumetinib were here investigated (29). Trametinib shows equal potency for targeting MEK1 and MEK2 and preferentially binds unphosphorylated MEK1/2 and thereby preventing Raf-dependent MEK phosphorylation and activation (30, 31). Selumetinib targets the unique inhibitor binding pocket adjacent to the Mg-ATP in MEK1/2. Sticking to this specific region causes a conformational change in unphosphorylated MEK1/2 resulting in a catalytically inactive position and blocking MEK1/2 from accessing the ERK1/2 activation loop. Selumetinib does not block binding and phosphorylation by Raf, which is different from trametinib (32). In addition, selumetinib shows higher potency to target MEK1 compared to MEK2. These different binding properties of selumetinib compared to trametinib result in higher IC50 for selumetinib in MEK sensitive tumors (reported IC50 values of 50 nM for trametinib and 2.5 μM for selumetinib using the A549 bronchioloalveolar carcinoma cell monolayer cultures (33)) which is in agreement with work done by Gilmartin et al. (30) and Yamaguchi et al. (34). The study of Fernandez et al. (35) marks differences in MEK efficacy in low-grade serous ovarian cancer cell lines as trametinib was found to be highly effective in blocking p-ERK1/2 compared to selumetinib (IC50 values were in the nM range for trametinib versus the μM-range for selumetinib). These findings are also in line with our observations regarding a different sensitivity for both MEK inhibitors with the established PM-LGSOC-01 cells. In vivo evaluation of trametinib in PM-LGSOC-01 revealed a similar sensitivity suggesting that the peritoneal stroma does not affect the trametinib response. Due to the failure of MEK inhibitors such as binimetinib in a phase 3 clinical trial for LGSOC and the unknown molecular mechanisms related to this failure (8), we strongly believe that the current model will assist in the better understanding of responsiveness and resistance to MEK inhibitors.

Establishing and analysing additional LGSOC lines might substantiate our finding and may provide a unique opportunity to study LGSOC progression and chemosensitivity.

## Additional Information

### Ethical approval and informed consent

Informed consent of the patients to use tumor material was obtained after the study protocol was approved by the institutional review board of the Ghent University Hospital. Animal experiments were conducted in accordance with the local ethics committee (ECD 15/28, Ghent University Hospital).

### Authors’ contributions

EDT acquired and interpreted *in vivo* and *in vitro* data, wrote the manuscript. KVdV and JVD performed pathological studies and interpreted pathological data. JVdM, JT, GW and GB acquired and interpreted *in vitro* and molecular data. HD and WC provided clinical interface, coordination and support. ODW coordinated the research project, designed experiments and wrote the manuscript. Critical revision of the manuscript and approval of the final version of the manuscript: all authors.

## Acknowledgments

The authors would like to thank S. Decloedt for the technical assistance and the preclinical core imaging facility of Ghent University (INFINITY) for providing the in vivo imaging systems. We are grateful to the people who consented to donate their tissues to support this work.

